# It only takes seconds for a human monoclonal autoantibody to inhibit N-methyl-D-aspartate receptors

**DOI:** 10.1101/2024.05.28.595700

**Authors:** Shang Yang, Johanna Heckmann, Abdulla Taha, Shiqiang Gao, Stephan Steinke, Michael Hust, Harald Prüß, Hiro Furukawa, Christian Geis, Manfred Heckmann, Jing Yu-Strzelczyk

**Affiliations:** Institute for Physiology, Department of Neurophysiology, Julius-Maximilians-Universität of Würzburg, 97070 Würzburg, Germany; Janelia Research Campus, Howard Hughes Medical Institute, 20147 Ashburn, United States; Jena University Hospital, Department of Neurology/Section of Translational Neuroimmunology, Am Klinikum 1, 07747 Jena, Germany; Technische Universita□t Braunschweig, Institut fu□r Biochemie, Biotechnologie und Bioinformatik, Department Medizinische Biotechnologie, Braunschweig 38106, Germany; Department of Neurology and Neuroscience, Autoimmune Neurology Group, Mayo Clinic, 32224 Jacksonville, USA; Department of Neurology and Experimental Neurology, Charité, Universitätsmedizin Berlin, 10117 Berlin, Germany; W.M. Keck Structural Biology Laboratory, Cold Spring Harbor Laboratory, Cold Spring Harbor, 11724 New York, USA

**Keywords:** Anti-NMDA receptor encephalitis, Patch-clamp recording, Autoantibody

## Abstract

Transfer of autoantibodies targeting ionotropic N-methyl-D-aspartate receptors in autoimmune encephalitis patients into mice leads to typical disease signs. Long-term effects of the pathogenic antibodies consist of immunoglobulin G-induced crosslinking and receptor internalization. We focused on the direct and immediate impact of a specific pathogenic patient-derived monoclonal autoantibody (immunoglobulin G #003-102) on receptor function.

We performed cell-attached recordings in cells transfected with the GluN1 and GluN2A subunit of the N-methyl-D-aspartate receptor. Immunoglobulin G #003-102 binds to the amino-terminal domain of the glycine-binding GluN1 subunit. It reduced simultaneous receptor openings significantly compared to controls at both low and high glutamate and glycine concentrations. Closer examination of our data in 50-second to 2-second intervals revealed, that Immunoglobulin G #003-102 rapidly decreases the number of open receptors. However, antigen-binding fragments of immunoglobulin G #003-102 did not reduce the receptor openings.

In conclusion, patient-derived immunoglobulin G #003-102 inhibits N-methyl-D-aspartate receptors rapidly and directly before receptor internalization occurs and the entire immunoglobulin G is necessary for this acute inhibitory effect. This suggests an application of the antigen-binding fragment-like constructs of #003-102 as a potential new treatment strategy for shielding the pathogenic epitopes on the N-methyl-D-aspartate receptors.

## Introduction

N-methyl-D-aspartate (NMDA) receptors are ionotropic glutamate receptors that play a central role in brain development and function^1,2^. Anti-NMDA receptor encephalitis is the most common cause of nonviral, autoimmune brain inflammation and is characterized by psychosis, dyskinesias, autonomic dysfunction, and seizures^3,4^. The clinical diagnosis relies on detecting NMDA receptor-binding antibodies in patients’ cerebral spinal fluid. Symptoms are believed to arise due to autoantibodies internalizing surface NMDA receptors in central neurons^5-7^. Most patients show a favourable treatment response upon immunotherapy^8^.

The NMDA receptors consist of two GluN1 subunits responsible for binding glycine and two glutamate-binding GluN2 or glycine-binding GluN3 subunits^1^. Three main domains make up each subunit^9^ (Fig 1): I) the amino-terminal domain (ATD), known for its role in allosteric modulation (e.g. through zinc and protons) and considered the primary epitope for autoantibody binding; II) the ligand binding domain (LBD), responsible for binding to both glutamate and glycine, as well as competitive antagonists; and III) the channel gate and pore are formed by the transmembrane domain (TMD)^10,11^. In addition to glycine and glutamate, proper activation of NMDA receptors hinges on relieving voltage-dependent channel blocks by Mg^2+^ ions^12^ by depolarization. The simultaneous ligand binding, Ca^2+^ influx, and depolarization lead to NMDA receptor-mediated synaptic plasticity, a cellular model for learning and memory^13^.

**Figure 1.**
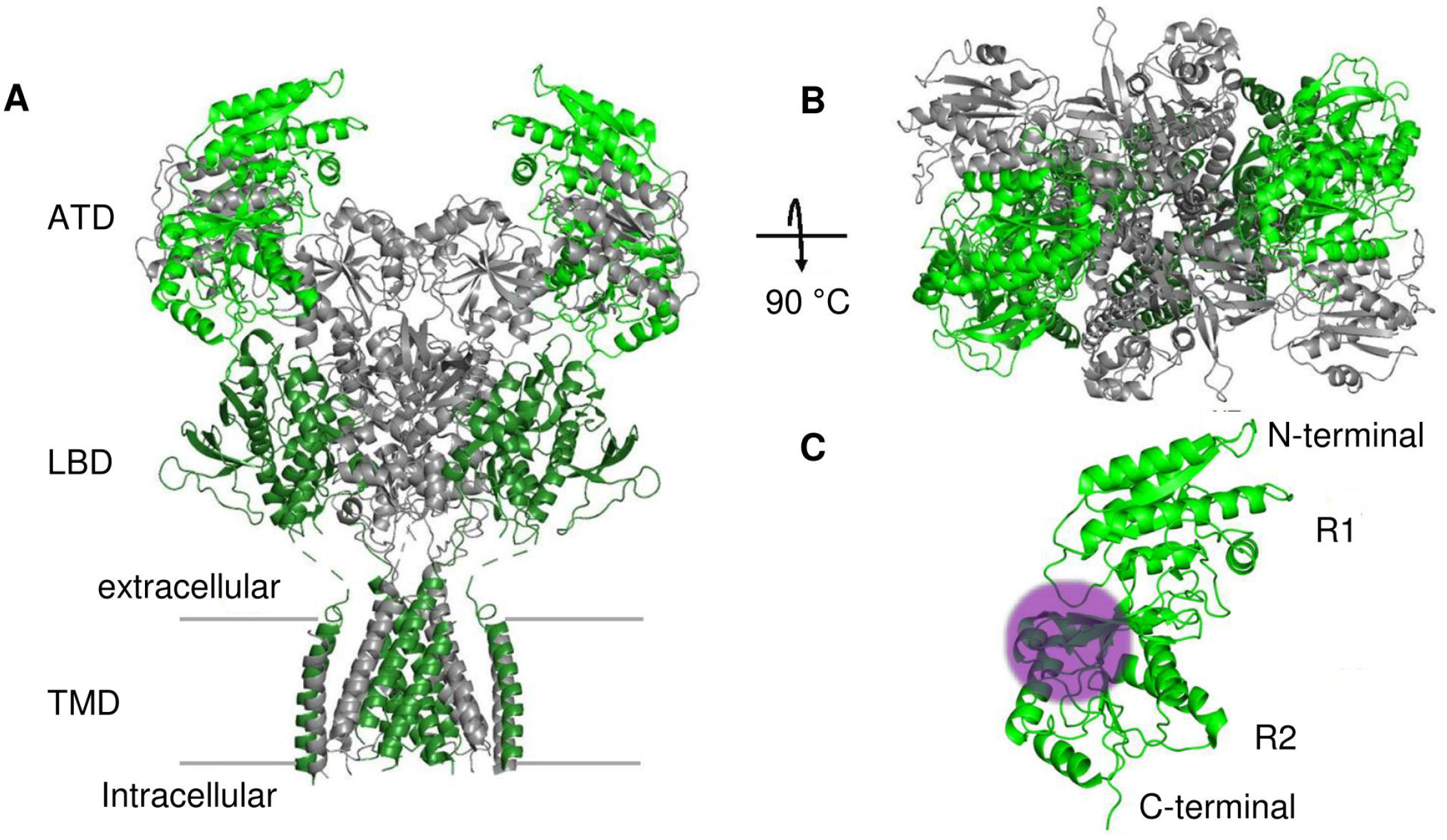
NMDA receptor structure with two GluN1 and GluN2A subunits. **(A)** Side view of the receptor in the 2-knuckle-sym conformation with GluN2A ATDs in open clam-shell conformation (in grey) and GluN1-subunits in green (PDB 6MMR)^9^. **(B)** Top view of the receptor. **(C)** Enlarged two lobes, R1 and R2, of the GluN1-ATD. The antibody may, e.g., bind to the top of the bottom lobe close to the hinge region between R1 and R2 of the extracellular ATD of the GluN1 subunit. Typical epitope dimensions are highlighted by an arbitrarily placed circular area with a diameter of 3 nm in magenta. The illustration was generated with PyMOL Molecular Graphics System, Version 2.0 Schrödinger, LLC.

Several GluN1-targeting antibodies were identified from the cerebrospinal fluid of anti-NMDA receptor encephalitis patients by isolating full-length immunoglobulin heavy and light chain genes through single-cell cloning of memory B cells and antibody-secreting cells^7^. The binding of monoclonal GluN1 antibodies to the ATD (Fig. 1C) is considered to result in crosslinking and internalization of NMDA receptors, which is presumed to be a critical factor in developing disease pathogenesis^7^.

Additionally, it is becoming increasingly evident that, in certain autoimmune disorders, even the binding of antibody fragments can alter receptor activity without causing internalization of the receptors. For example, in another form of autoimmune encephalitis where autoantibodies attack the GABA_A_ receptor, the administration of antigen-binding fragments (Fabs) has been observed to impede the activity of the inhibitory ionotropic receptor^14^. Moreover, in a particular type of Graves’ disease, which is the primary cause of hypothyroidism unrelated to iodine deficiency, autoantibodies mimicking hormone action at the thyrotropin receptor trigger inappropriate receptor activation^15,16^.

Furthermore, after immunization of mice with NMDA receptors, a murine antibody was identified that specifically binds to the GluN1-GluN2B NMDAR subtype and down-regulates ion channel function allosterically^17^. On the other hand, autoantibodies from patients with Lupus erythematosus can enhance NMDA receptor activity by acting as positive allosteric modulators for GluN2A^18^. This indicates that antibody binding can directly modulate NMDA receptor function. The diverse and complex action modes of anti-NMDA receptor autoantibodies were recently reviewed^2^. Early patch-clamp recordings revealed that acute exposure to polyclonal IgG antibodies from patients with NMDAR encephalitis prolongs the receptor open time in anti-NMDA receptor encephalitis^19^. Against this, glutamate-evoked currents in *Xenopus* oocytes were more recently found to be reduced upon the application of patient-derived NMDA receptor antibodies^20^. Because of these discrepant results, it is important to explore direct and acute effects of antibodies from patients with anti-NMDA receptor encephalitis in more detail.

Therefore, we tested for acute effects of a patient-derived and well-characterized pathogenic monoclonal anti-GluN1 antibody, Immunoglobulin G (IgG) #003-102, and its Fab fragment on GluN1/GluN2A NMDA receptors using cell-attached single channel patch clamp recordings. We find that IgG #003-102, but not its Fab fragments, rapidly reduces the number of simultaneously open NMDA receptors at low and high agonist concentrations.

## Materials and methods

Full details of the methods are provided in the Supplementary material.

## Results

### Cell-attached patch clamp recording

The GluN1 binding monoclonal IgG #003-102 was identified from an antibody-secreting cell of an 18-year-old female patient with ovarian carcinoma one week after the onset of acute encephalitis^7^. Our approach to test for direct effects of antibody binding to the receptor was unbiased and highly sensitive. Patch clamp measurements allow different recording configurations^21^. Cell-attached recording isolates approximately a 1 µm^2^ small membrane area and enables the recording of the activity of single receptors. In addition, it requires a minimal number of antibodies, which is applied only to the solution in the recording electrode acutely and at a defined concentration. We performed blinded experiments to compare the acute effects of control IgG mGO53, IgG #003-102, Fab fragment of IgG mGO53, and Fab fragment of IgG #003-102 on NMDA receptor opening using cell-attached patch-clamp recordings (Supplementary Fig. 1). Antibodies and Fab fragments were introduced via number-coded vials and applied to the pipette solution (Supplementary Fig. 1B), which reached the extracellular domain of the NMDA receptor (Supplementary Fig. 1C). Control IgG (mGO53) or patient IgG #003-102 were used in a saturating concentration of 10 μg/ml since the binding constant of IgG #003-102 to NMDA receptors is 1.16 μg/ml^22^. YFP-fluorescent HEK cells were selected for recording (Supplementary Fig. 1D). Once a seal was formed, the final antibody concentration was reached at the receptors. After completing experiments and trace analysis, the experimenter received the key to the number codes and evaluated the statistics.

### Acute effect of IgG #003-102 at low agonist concentrations

Representative 500 ms long original traces of cell-attached recordings are shown in Fig. 2A. Fig. 2A exhibits downward current steps with up to four simultaneous open receptors with control IgG. IgG #003-102 was used in the recording below, and no more than two receptors opened simultaneously. In a total of 52 recordings of this type with receptor openings at low agonist concentrations around the EC50 (4 µM glutamate, 1µM glycine), we observed one to eight simultaneous receptor openings in controls (n = 29, median = 4) and one to seven openings with IgG #003-102 treatment (n = 23, median = 3). IgG #003-102 significantly reduced the number of simultaneously open NMDA receptors (Fig. 2B, *P* = 0.033). The histograms in Fig 2B illustrate that we obtained putative “single” channel recordings in only five control recordings and 9 with patient IgG.

**Figure 2.**
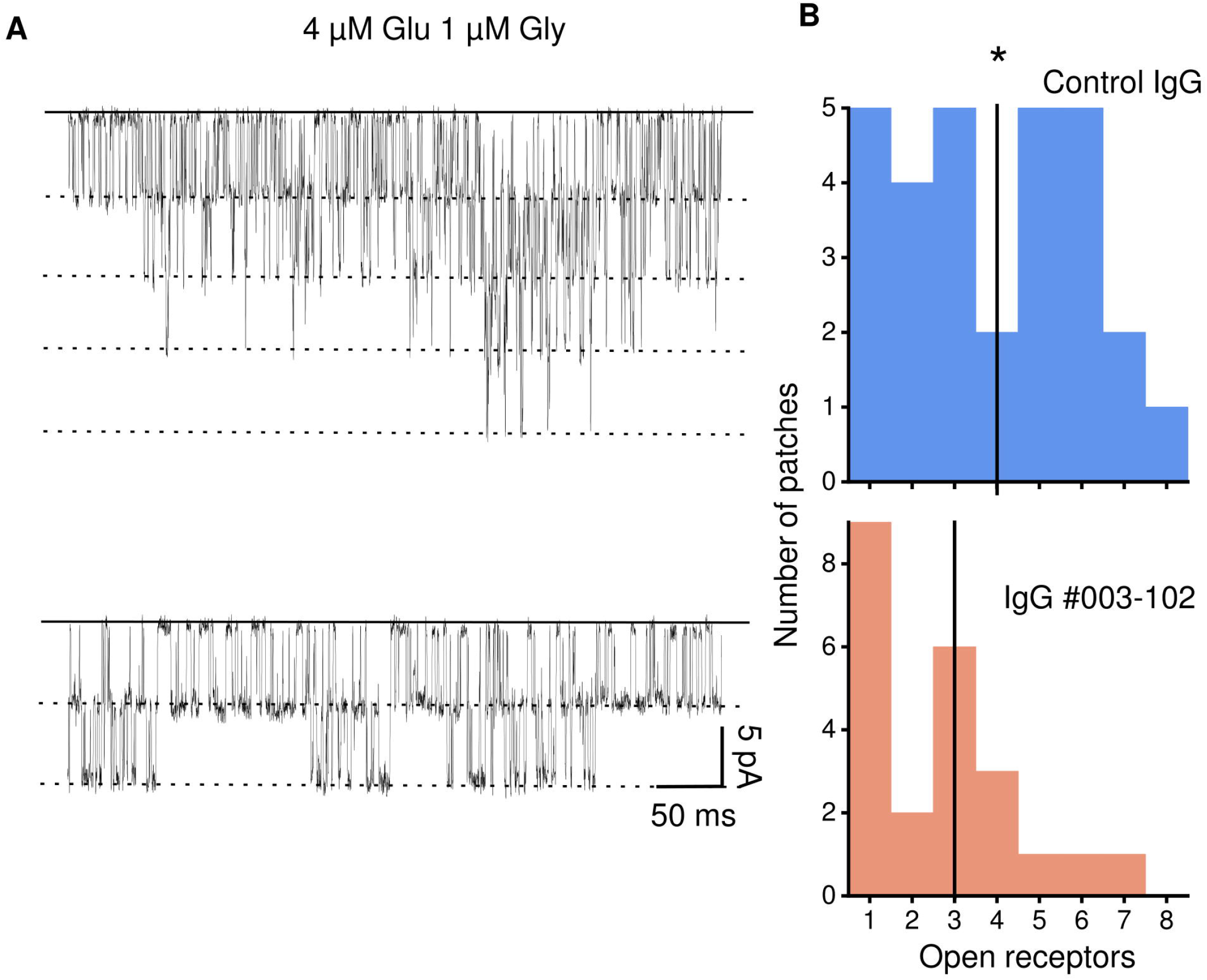
Acute effects of patient-derived monoclonal IgG #003-102 on NMDA receptor opening at low agonist concentrations. Fifty-two recordings were performed and evaluated in a blinded fashion with either control IgG (mGO53) or patient IgG #003-102 in the concentration of 10 μg/ml with 4 μM glutamate and 1 μM glycine. (A) Representative recording with control IgG and up to four simultaneously open NMDA receptors (upper panel) and representative recording with patient IgG #003-102 and up to two simultaneously open NMDA receptors (lower panel). (B) Histogram of the number of patches with one to eight simultaneously open NMDA receptors in controls (upper panel in light blue, n = 29 patches, median = 4, indicated by the black line). Histogram of the number of patches with one to seven simultaneously open NMDA receptors with patient IgG (003-102, lower panel in light red, n = 23 patches, median = 2.5, indicated by black line). A Mann-Whitney rank-sum test gives a significant *P* = 0.033 for comparing the control and patient IgG #003-102 results.

### Acute effects of IgG #003-102 at high agonist concentration

The receptor structure is known to be agonist-dependent, and it thus appeared attractive to test the effects of antibodies at higher concentrations^1^. Like the result at low agonist concentrations, at high agonist concentrations (1 mM glutamate, 0.1 mM glycine), the number of simultaneously open NMDA receptors decreased dramatically upon exposure of IgG #003-102 (Fig. 3A, 38 control recordings, median = 3; 62 IgG #003-102 recordings, median = 2, *P* = 0.002). However, despite a one hundred-fold or more difference in glutamate and glycine concentration in the recordings for Fig. 2B and 3A, the number of simultaneously open receptors was similar for the two groups of control and IgG #003-102 treated receptors. Strikingly, we obtained putative “single” channel recordings in only 9 of our 38 recordings with control antibody at high glutamate and glycine concentration compared to 29 patches out of 62 with patient antibody (Fig. 3A).

**Figure 3.**
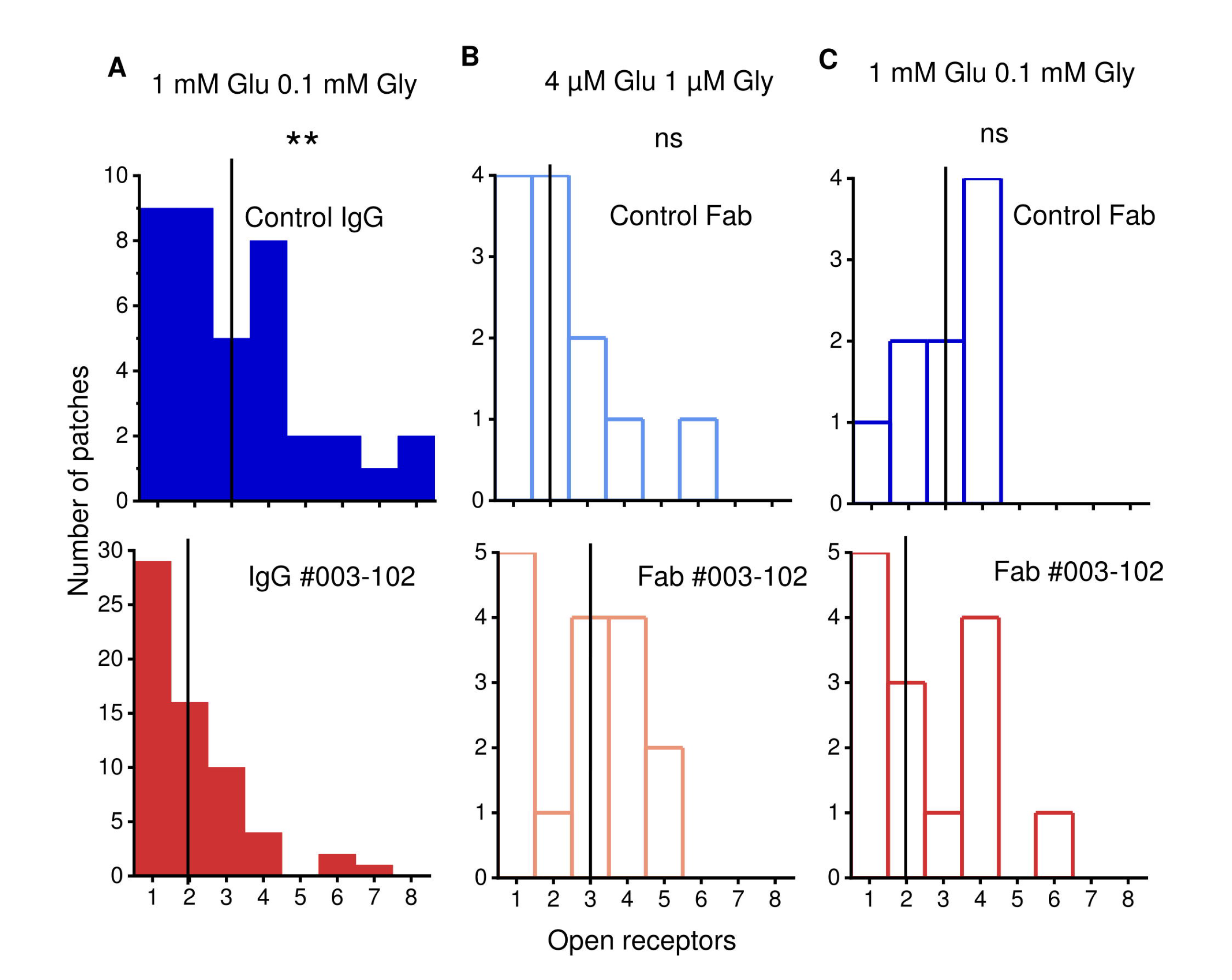
Acute effects of IgG #003-102 and Fabs from IgG #003-102 on NMDA receptor openings. (A) One hundred recordings were performed and evaluated blinded with 1 mM glutamate and 0.1 mM glycine and either control IgG (mGO53) or patient IgG #003-102 in a 10 μg/ml concentration. The upper panel in blue is a histogram of the number of patches with one to eight simultaneously open NMDA receptors in controls (n = 38 patches, median = 3, indicated by a black line). The lower panel in red: Histogram of the number of patches with one to eight simultaneously open NMDA receptors with monoclonal patient IgG #003-102 (n = 62 patches, median = 2, indicated by the black line). A Mann-Whitney rank-sum test gives a significant *P* = 0.002 for comparing the control and patient IgG #003-102 results. (B) Histogram of the number of patches with one to six or one to five simultaneously open NMDA receptors in recordings with 4 μM glutamate plus μM glycine and control Fabs (upper panel in light blue, n = 12 patches, median = 2) or Fabs of IgG #003-102 in a concentration of 7 μg/ml (lower panel in light red, n = 16 patches, median = 3, *P* = 0.35). (C) Histogram of the number of patches with one to four or one to six simultaneously open NMDA receptors in recordings with one mM glutamate and 0.1 mM glycine and control Fabs (upper panel in blue, n = 9 patches, median = 3) and patches with Fabs of IgG (003-102) in a concentration of 7 μg/ml (lower panel in red, n = 14 patches, median = 2, *P* = 0.4).

### No effect of Fab antibody fragments on NMDA receptor opening

Given the inhibitory effect of IgG #003-102, we were curious to test if Fab antibody fragments would also be effective, as recently found in another form of autoimmune encephalitis and antibodies to the GABA_A_ receptor^14^. The histograms in Fig. 3B and C show data from experiments with either control Fabs or Fabs of IgG #003-102 at a concentration of 6.67 µg/mL and either low or high agonist concentrations. These experiments revealed that Fabs of IgG #003-102 do not downregulate ion channel activity (at low agonist concentration, 12 recordings with control Fabs, median = 2, 16 recordings with IgG #003-102, median = 3; *P* = 0.35; at high agonist concentration, nine recordings with control Fabs, median = 3, 14 recordings with IgG #003-102, median = 2; *P* = 0.4).

### NMDA receptor openings are inhibited within seconds by patient-derived monoclonal IgG #003-102

To determine the time course of the inhibitory effect, we analyzed the number of open receptors in each recording at high agonist concentration in 50 s segments. In the first 50s, the inhibitory effect was already significant. We further analyzed the first 50 s of the recordings in 10 s segments and found that the inhibition is already apparent even in the first 10 s of our recordings. Further subdivision into 2 s windows revealed an immediate inhibition even at this temporal resolution (Fig.4).

### The kinetics of the remaining NMDA receptors are not altered

To find out if the kinetics of the NMDA receptor would be altered by IgG #003-102, we analyzed the recordings with only one active channel in the patch (Supplementary Fig. 2). The open time and closed time and their distributions are similar in both conditions (Supplementary Fig. 2A and B). Although NMDA receptor opening is known to be quite variable in NMDA receptors and shifts from one mode to another^23^, the mean open time was relatively stable throughout our recordings.

## Discussion

Our data reveal that a pathogenic patient-derived monoclonal anti-NMDA receptor antibody, but not its Fab fragment, acutely reduces NMDA receptor openings within seconds. We used the patient IgG antibodies and Fabs at such high concentrations in our electrodes that receptor binding should have happened within less than a minute upon forming a patch^22^. Due to the short time course of the antibody application in our cell-attached recordings, receptor internalization can be excluded as the underlying mechanism for the reduction of receptor openings as NMDA receptors are relatively immobile membrane proteins, and receptor internalization is too slow to be relevant compared to our 150-second-long recordings^24,25^.

We used cell-attached recording, which allowed us to record currents from individual open receptors. Blind experiments ensured unbiased evaluation and a large dataset showed significant differences when comparing control and patient antibodies. We obtained data from patches with up to 8 open NMDA receptors, and one could argue that it is favourable when testing for functional effects of a compound to have many channels in a patch. Two or more receptors opened simultaneously in a substantial fraction of our patches (Fig. 2 and Fig 3). Therefore, we used the peak number of simultaneously open receptors as the lower limit of the number of functional receptors per patch^26,27^. We used this simple measure to assess the effects of the different applications. An advantage of our approach is that it does not require any data selection. The temporal evaluations for Fig. 4 should be considered with some caution. Electrode solution is usually blown out of the electrode tip before patch formation. Thus, antibodies reached the receptors, albeit at a much lower concentration, even before patch formation. Furthermore, it usually takes additional seconds before recordings start after obtaining a seal (since the test pulse needed to be turned off, the desired holding potential turned on, and data acquisition also turned on). If seal formation was not immediate, further delays occurred. Collectively, Fig. 4 thus only allows us to conclude that the inhibition happened within seconds. The rate of antibody binding needs to approach the diffusion limit of about 10^7^ M^-1^ s^-1^ for IgG for such a rapid inhibition. It would be worthwhile to investigate this further for bivalent and monovalent IgG #003-102. In outside-out patches, another patch clamp configuration^21^ combined with a piezo-driven system antibody applications should be achievable within less than a millisecond. However, this setup requires large amounts of antibodies, and the rundown of NMDA receptor currents in outside-out patches will make it difficult to further probe the kinetics of patient antibodies inhibit with this approach.

**Figure 4.**
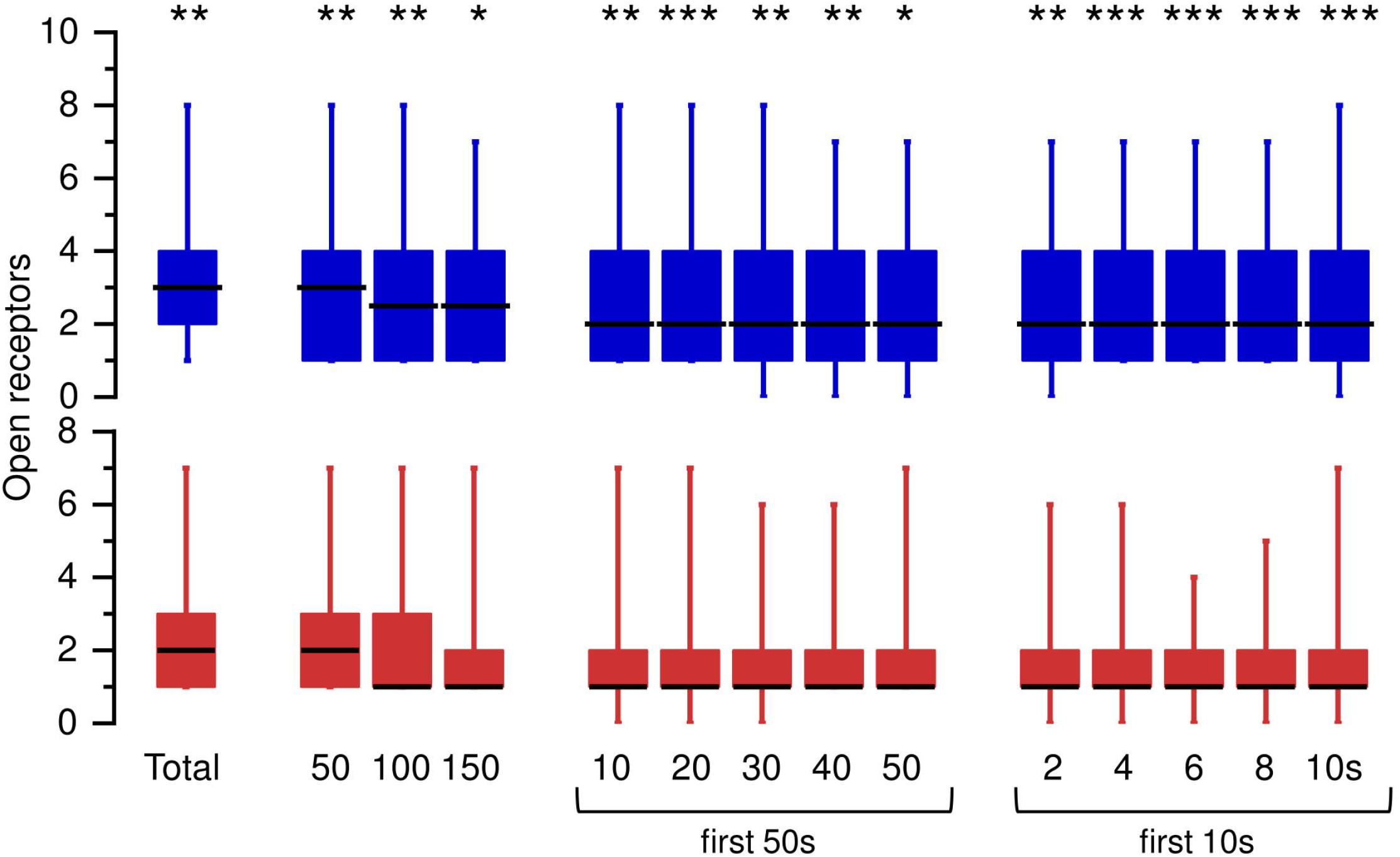
IgG #003-102 inhibits NMDA receptors within seconds. The results for all recordings in Fig. 3A are shown as total on the left. The following three entries (50, 100, 150) show evaluations of this data split up into three 50-second segments. The first 50 seconds of the recordings were then further analyzed in five 10-second segments (10, 20, 30, 40, 50), and the first 10 seconds were again divided into 2-second segments (2, 4, 6, 8, 10) as indicated below the data plots. Significance levels are given as: * *P* < 0.05, ** *P* < 0.01, *** *P* < 0.001.

In anti-NMDA receptor encephalitis, antibodies bind to the ATD of the GluN1. Although the ATD does not directly interact with glutamate or glycine, it plays a crucial role in unlocking the receptor for activation^28^. Zinc and protons are physiologic allosteric modulators of NMDA receptors and bind to the ATD^9^. Antibody effects might differ depending on the exact site of the epitope on the ATD and glycine, zinc, and proton concentrations during recordings. In our recordings, EDTA was used to minimize gating effects of the divalents.

Further investigations at lower pH and with higher extracellular zinc concentration are necessary to cover the whole physiologic range of NMDA receptor function. The effects described here might be more or less pronounced under these conditions. In addition, it appears desirable to unite structural and functional insights regarding the direct and acute effects of specific human autoantibodies on NMDA receptor function, similar to recent work on murine antibodies^17^. Furthermore, other antibodies against the ATD of NR1 should be tested, and it appears worthwhile to test for the interaction of different antibodies in the ATB and shielding of epitopes.

Our recordings revealed that receptor kinetics are not influenced by the antibodies (Supplementary Fig. 2). The normal receptor kinetics and a reduction of simultaneously open receptors with patient-derived antibodies lead to the conclusion that patient-derived antibodies bind and lock receptors in long-lived closed states acutely. In contrast, the rest of the receptors are unaffected. This interpretation aligns with the inhibitory effect of murine antibodies, which recognize the ATD of the GluN2B subunit and increase the population of the non-active state ^17^. However, in contrast to our findings, in Tajima et al., 2022, Fabs were also effective in downregulating GluN1-GluN2B NMDAR activity^17^. Furthermore, in anti-GABA_A_ receptor encephalitis, GABA receptor antibodies and their Fabs inhibit the GABA-binding site and lock the receptor in a resting-like state^14^. In our case, the binding of one individual epitope on the NMDA receptor is insufficient to inhibit the GluN1-GluN2A NMDA receptor.

Immunotherapy is the standard of care for anti-NMDA receptor encephalitis so far. New therapeutic approaches aim to neutralize the pathogenic antibodies before they can bind to the receptor^29^. Based on our results, we propose the design of an engineered monovalent IgG as an additional new therapeutic approach that should neither interfere with receptor function nor induce receptor internalization. Our data shows that IgG #003-102 needs to be bivalent for the acute inhibitory effect of the NMDA receptor, as the Fabs do not inhibit the receptors. The limiting factor here should be the distance between the two IgG paratopes, which could be tested by engineering IgGs with flexible linkers to determine an optimal paratope distance. Moreover, it is known that the IgG paratopes need to crosslink the NMDA receptors to induce internalization^5^. Therefore, whether monovalent IgG #003-102, e.g., by using a heterodimeric IgG with one heavy chain without a Fab and one heavy chain with the Fab part, or by using the corresponding Fab reduces symptoms by preventing both acute receptor inhibition and internalization should be tested in an *in vivo* model.

Positive allosteric modulators of NMDA receptors effectively upregulate the activation and charge transfer of the remaining NMDA receptors with antibodies from patients with anti-NMDA encephalitis^30^. It would also be interesting to test whether positive allosteric modulators of NMDA receptors can compensate for the acute inhibitory effects on NMDA receptor currents described here. In summary, we show that upon antibody binding, an acute receptor inhibition occurs in addition to later receptor crosslinking and internalization. These mechanisms might contribute to disease pathology in anti-NMDA receptor encephalitis.

## Acknowledgments

This section is not mandatory.

## Supplementary material

Supplementary material is available at Brain online.

## Abbreviations

ATD: Amino-Terminal Domain
DMEM: Dulbecco’s Modified Eagle Medium
EGTA: Ethyleneglycol-bis(β-aminoethyl)-N,N,NC,NC-tetraacetic Acid
Fab: Fragment Antigen-Binding
HEK cells: Human Embryonic Kidney cells
IgG: Immunoglobulin G
LED: Light-Emitting Diode
LBD: Ligand Binding Domain
NMDA: N-methyl-D-aspartate
TMD: TransMembrane Domain
YFP: Yellow Fluorescent Protein

