## Supplemental Material for "It only takes seconds for a human monoclonal autoantibody to inhibit N-methyl-D-aspartate receptors"

# to

### Materials and methods

#### Cloning

The rat GluN1 and GluN2A DNA segments were provided by Prof. Noam Ziv (Technion, Haifa, Israel) and Dr. Georg Köhr (Central Institute of Mental Health, Mannheim, Germany), and were inserted into the pcDNA3.1 vector within the N-terminal BamHI and the C-terminal HindIII restriction sites.

#### Cell culture and transfection

Human embryonic kidney cell-line (HEK293T) cells were cultured at 37°C and 5% $CO_2$ in Dulbecco's modified Eagle medium (DMEM) supplemented with 10% fetal bovine serum. The cells were seeded in 24-well plates (Thermo Scientific™ Nunc™ 24-well plates, Ref:142475) on poly-lysine coated coverslips (round, Ø 10 mm) with 500 µl medium. Transfection was performed 24 hours after seeding using Lipofectamin™ 3000 transfection reagent. For the visual selection of transfected cells, 500 ng soluble yellow fluorescence protein (YFP) in pcDNA3.1 was transfected with 300 ng GluN1 and 300 ng GluN2A plasmids into HEK293T cells. 25 µl DMEM with 1µl Lipofectamine™ 3000 reagent and 25 µl DMEM with plasmids and 1µl P3000™ 3000 reagent were mixed well and incubated for 15 minutes at room temperature. This preparation is for the transfection of 4 wells and was completed by adding ketamine (1.26 mM) and (2R)-amino-5-phosphonovaleric acid (0.1 mM) to reduce excitotoxicity during culturing.

#### Production of IgG #003-102 as IgG and Fab

Gene strings for the heavy (VH) and light (VL) chains of antibody #003-102 (WO 2017/029299 AI) were synthesized and ordered from Gene Art (Thermo Fisher Scientific). The VH gene was cloned into the human IgG1 format using the pCSEH1c.2 vector and into the Fab format using the pCSE2.5-Hc-hFab.2 vector. The VL gene was subcloned into the pCSL3hk.2-Xp vector. Both cloning procedures were adapted for Golden Gate Assembly with the Esp3I restriction enzyme (New England Biolabs, Frankfurt, Germany). The Golden Gate cloning methodology followed the protocol previously described by Steinke et al[1]. Antibody production and purification protocols were conducted according to Bertoglio et al[2] and Steinke et al[1]. Expi293F cells were cultured at 37°C with 5% $CO_2$ and 110 rpm in Gibco FreeStyle F17 expression media (Thermo Fisher Scientific). The media was supplemented with 8 mM Glutamine and 0.1% Pluronic F68 (PAN Biotech). For transfection, DNA vectors for IgG production were mixed in a 1:1 ratio and combined with 40 kDa polyethyleneimine (PEI) (Polysciences). The mixture was incubated for 25 minutes at room temperature before being added to the cells. After 48 hours, the culture volume was doubled by adding HyClone SFM4Transfx-293 media (GE Healthcare) supplemented with 8 mM Glutamine. Additionally, the HyClone Boost 6 supplement (GE Healthcare) was added at 10% of the final volume. One week post-transfection, the supernatant containing the antibodies was harvested by centrifugation at 1500 xg for 15 minutes. Antibodies were then purified using the Profinia System (BIO-RAD) or ÄKTA System (Cytiva Life Sciences). The IgG1 antibody format was purified using Protein A, while the Fab format was purified using a 6xHis-tag.

#### Cell-attached patch clamp recordings

Patch pipettes were manufactured with a DMZ-Puller from borosilicate glass (outer Ø 2.00 mm, inner Ø 1.16 mm, GB200F-10, Science Products) and had resistances between 10 - 20 MΩ. To reduce the capacitive noise, the patch pipette was filled with the pipette solution only at the very tip part (< 1 mm). Cells were visualized using an inverted microscope (Axiovert 30, Zeiss). Transfected cells were selected by identifying the YFP fluorescence with an excitation 490-510 nm, dichroic mirror 515 nm, and emission 520 -550 nm filter cube. The measurements were performed in the cell-attached configuration[3] at 23°C. Recordings for Figure 2 were performed with identical pipettes and bath solutions containing 150 mM NaCl, 10 mM HEPES, and 0.05 mM EDTA at pH 8. In Figure 5, D-PBS (Gibco, 138 mM NaCl, 8.1 mM $Na_2HPO_4$, 2.67 mM KCl, and 1.47 mM $KH_2PO_4$) was used in a bath, and 150 mM NaCl, 2.5 mM KCl, 10 mM HEPBS, 1 mM EDTA, pH 8 with NaOH as pipette solution. Figures 3 and 4 show results from both conditions. The channel opening probability was maximized using the calcium chelator (which minimizes gating effects of divalents)[4] and a basic pH (since protons also reduce channel opening)[5]. Before the electrophysiological experiment, glycine (1µM or 0.1 mM) and glutamate (4µM or 1 mM) and either control IgG (mGO53) or #003-102 at a concentration of 10 µg/mL or their Fabs at a concentration of 6.7 µg/mL were applied to the pipette solution in a blinded fashion as illustrated in Supp. Fig. 1. Recordings were done with an Axopatch 200B amplifier (Axon Instruments INC.), a power1401-3A interface (Cambridge Electronic Design Limited), and Spike2 version 7 (Cambridge Electronic Design Limited) for data acquisition.

#### Data analysis

The traces were sampled at 50 kHz, pre-filtered at 10 kHz, and 2 kHz (Bessel) for evaluation and display. Recordings with unstable baselines or amplitudes were not used for further analysis. Clampfit 10.7 was used for event detection and determining the open probability of recordings with only one channel in the patch.

We estimated the number of functional channels in the patch by counting the maximum number of simultaneously open channels[6,7]. Origin Pro 2022 was used for statistics. Mann-Whitney rank-sum test and students' t-tests were used to determine significance.

#### Data availability

The datasets produced in this study are available upon request.

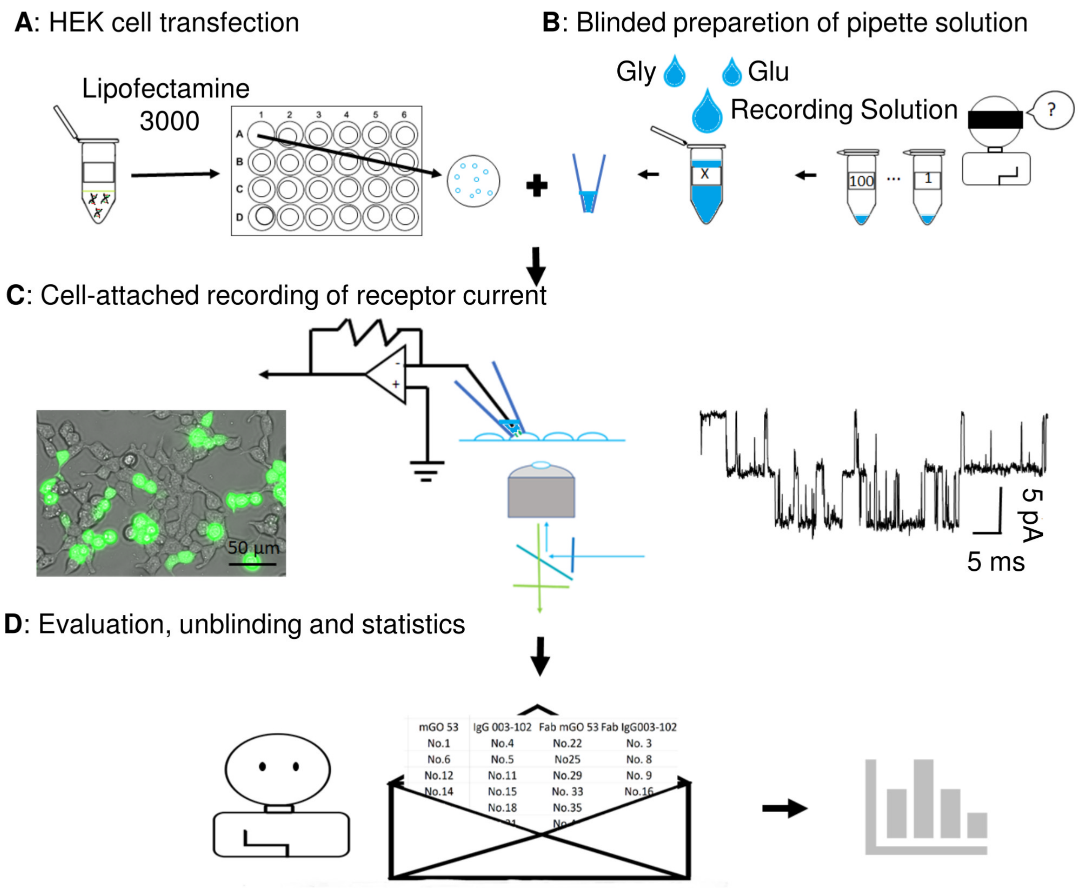


**Supplementary Figure 1. Experimentally examining the short-term effects of monoclonal patient autoantibodies on NMDA receptors. (A)** Cells were seeded on round coverslips and transfected with NMDA receptor and YFP plasmids. **(B)** Aliquots of either mGO53 (control IgG), patient IgG #003-102, Fab fragment of mGO53 (control fab), or Fab fragment of IgG #003-102 were added to numbered test tubes. In a blinded fashion, the experimenter received number-coded tubes. The pipette solution was prepared by adding 1 ml recording solution, glutamate, and glycine to reach the final concentration of 10 µg/ml for IgGs and 6.7 µg/ml for fabs, respectively. (C) YFP-expressing cells were selected for cell-attached recording. (D) After completing the recordings and the data analysis, experimenters were unblinded, received the key of the number codes, and evaluated the data.

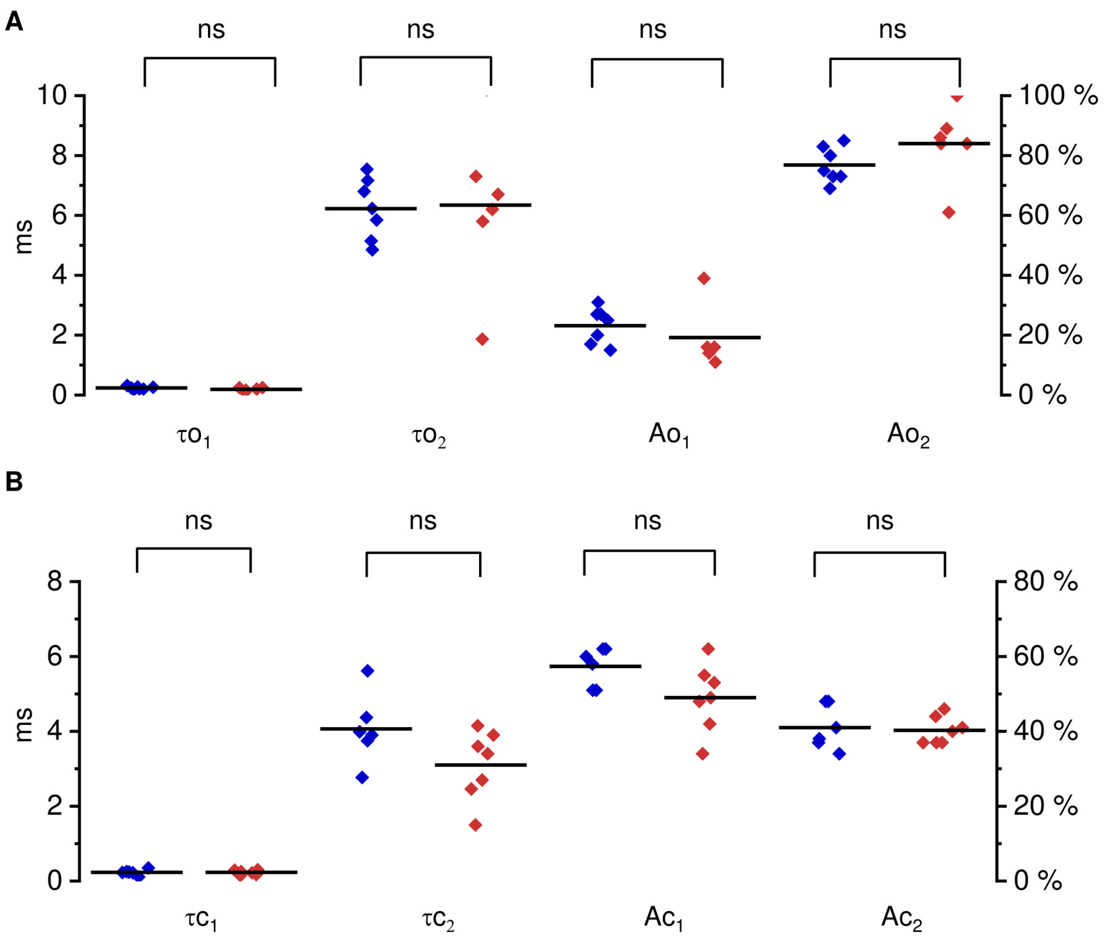


**Supplementary Figure 2. IgG #003-102 alters neither the open time nor the closed time of the NMDA receptors.**

**(A)** The open time of NMDA receptors in control recordings (mGO53, n = 7) was fitted with a mixture of two exponentials with mean time constants of 0.24 ms ($\tau o_1$ relative area 23 %) and $\tau o_2$ = 6.2 ms (relative area 77 %), which are not significantly different from recordings with patient-derived IgG (IgG #003-102, mean $\tau o_1$ = 0.19 ms, relative area = 19 %, mean $\tau o_2$ = 6.3 ms, relative area = 81 %, n = 6). **(B)** The closed time of NMDA receptors in control recordings (n = 6) was also fitted with a mixture of two exponentials with mean time constants $\tau c_1$ of 0.23 ms (relative area 57%) and $\tau c_2$ 4.1 ms (relative area 49 %), which are again not significantly different from recordings with patient-derived IgG (mean $\tau c_1$ = 0.23 ms, relative area = 49 %, $\tau c_2$ = 4.0 ms, relative area = 31 %, n = 7).
